# Sustained TREM2 stabilization accelerates microglia heterogeneity and Aβ pathology in a mouse model of Alzheimer’s disease

**DOI:** 10.1101/2021.06.23.449405

**Authors:** Rahul Dhandapani, Marilisa Neri, Mario Bernhard, Irena Brzak, Tatjana Schweizer, Stefan Rudin, Stefanie Joller, Ramon Berth, Annick Waldt, Rachel Cuttat, Ulrike Naumann, Caroline Gubser Keller, Guglielmo Roma, Dominik Feuerbach, Derya R. Shimshek, Ulf Neumann, Fabrizio Gasparini, Ivan Galimberti

**Affiliations:** Neuroscience, Novartis Institutes for Biomedical Research, Basel, Switzerland; Chemical Biology and Therapeutics, Novartis Institutes for Biomedical Research, Basel, Switzerland; Roche Glycart AG, Schlieren, Switzerland

## Abstract

TREM2 is a transmembrane protein expressed exclusively in microglia in the brain that regulates inflammatory responses to pathological conditions. Proteolytic cleavage of membrane TREM2 affects microglial function and is associated with Alzheimer’s disease, but the consequence of reduced TREM2 proteolytic cleavage has not been determined. We generated a transgenic mouse model of reduced TREM2 shedding (Trem2-IPD) through amino acid substitution of ADAM-protease recognition site. We found that Trem2-IPD mice displayed increased TREM2 cell surface receptor load, survival and function in myeloid cells. Using single cell transcriptomic profiling of mouse cortex we show that sustained TREM2 stabilization induces a shift of fate in microglial maturation and accelerates microglial responses to Aβ pathology in a mouse model of Alzheimer’s disease. Our data indicate that reduction of TREM2 proteolytic cleavage aggravates neuroinflammation during the course of AD pathology suggesting that TREM2 shedding is a critical regulator of microglial activity in pathological states.

## Introduction

Alzheimer’s disease (AD) is a slow progressive neurodegenerative disease characterized by the deposition of aggregating proteins, chronic inflammation, neuronal loss and subsequent dementia (Holtzman et al., 2011, Bartels et al., 2020). Recent advancements in genome wide association studies (GWAS) have established an important role for genetic variants of microglial genes in the progression of AD (Jonsson et al., 2013, Bohlen et al., 2019, Guerreiro and Hardy, 2014, Sierksma et al., 2020). Of particular interest is TREM2, a transmembrane receptor expressed in cells of myeloid lineage including macrophages, microglia, dendritic cells and osteoclasts. In the brain, microglia activation via TREM2 signaling promotes cell survival, chemotaxis to the site of insult, induces expression of proteins involved in lipid metabolism and suppresses inflammation (Kleinberger et al., 2014, Wang et al., 2015, Turnbull et al., 2006, Ulland et al., 2017).

Under pathological conditions TREM2 undergoes enhanced ectodomain shedding by ADAM10/17 proteases resulting in release of soluble TREM2 (sTREM2) into cerebrospinal fluid (CSF) (Suárez-Calvet et al., 2016, Heslegrave et al., 2016, Piccio et al., 2016). TREM2 shedding is enhanced in the H157Y variant establishing a crucial link between TREM2 shedding and AD progression (Thornton et al., 2017). Nonetheless, how reduced TREM2 proteolytic cleavage affects the progression of AD remains unclear. Here, we sought to reduce proteolytic cleavage of TREM2 by genetically modifying the ADAM10/17 binding site on the stalk region of the receptor (Schlepckow et al., 2017, Feuerbach et al., 2017) thereby increasing cell surface load and enhancing TREM2 dependent microglial functions. We then used an APP23xPS45 mouse model of amyloidosis to investigate the effects of TREM2 stabilization on Aβ deposition and microglial gene signatures and to understand the therapeutic modality for sustained TREM2 stabilization in Alzheimer’s disease.

## Results

### Reducing membrane TREM2 cleavage by amino acid substitution increases microglial function

To engineer cleavage reduction of TREM2 we identified Ile-Pro-Asp (IPD) as the least preferred amino acids for the sub-site binding of ADAM10/17 based on previously published studies (Liu et al., 2014, Tucher et al., 2014) and generated transgenic mice constitutively harboring the IPD-variant (hereby called Trem2-IPD) in place of the native (His-Ser-Thr) sequence (Figure 1A). To establish the functional effects of the Trem2-IPD mice, we first measured membrane TREM2 levels in bone marrow derived macrophages (BMDM) by FACS. Under basal state, BMDM from Trem2-IPD mice showed higher TREM2 levels compared to WT (Figure 1B). Near similar levels of TREM2 were achieved upon treatment of BMDM with ADAM17 inhibitor, DPC333. Treatment with PMA on the other hand resulted in complete reduction of membrane TREM2 levels in WT BMDM. However, only a small reduction was observed in Trem2-IPD BMDM (Figure 1C). This indicates that modified TREM2 still undergoes physiological processing, albeit lower than WT. We then used acute brain slice cultures to induce brain injury and TREM2 activation in an ex-vivo preparation. Measurement of TREM2 detected in the culture medium revealed a time dependent increase of soluble TREM2 in the supernatant across both genotypes. However, culture media derived from Trem2-IPD slices showed lower sTREM2 levels as compared to WT slices (Figure 1D). We then tested the survival of BMDM upon M-CSF withdrawal by quantifying total ATP within the cells using a CellTiter Glo assay and observed enhanced survival in Trem2-IPD BMDM compared to WT BMDM in the absence of M-CSF (Figure S1A). Moreover, primary microglia from Trem2-IPD mice showed increased uptake of pHrodo-labelled bioparticles compared to WT primary microglia (Figure S1B-S1E). These results suggest that myeloid cells from Trem2-IPD mice show higher membrane TREM2, reduced TREM2 shedding, pro-survival and enhanced phagocytic activity.

**Figure 1:**
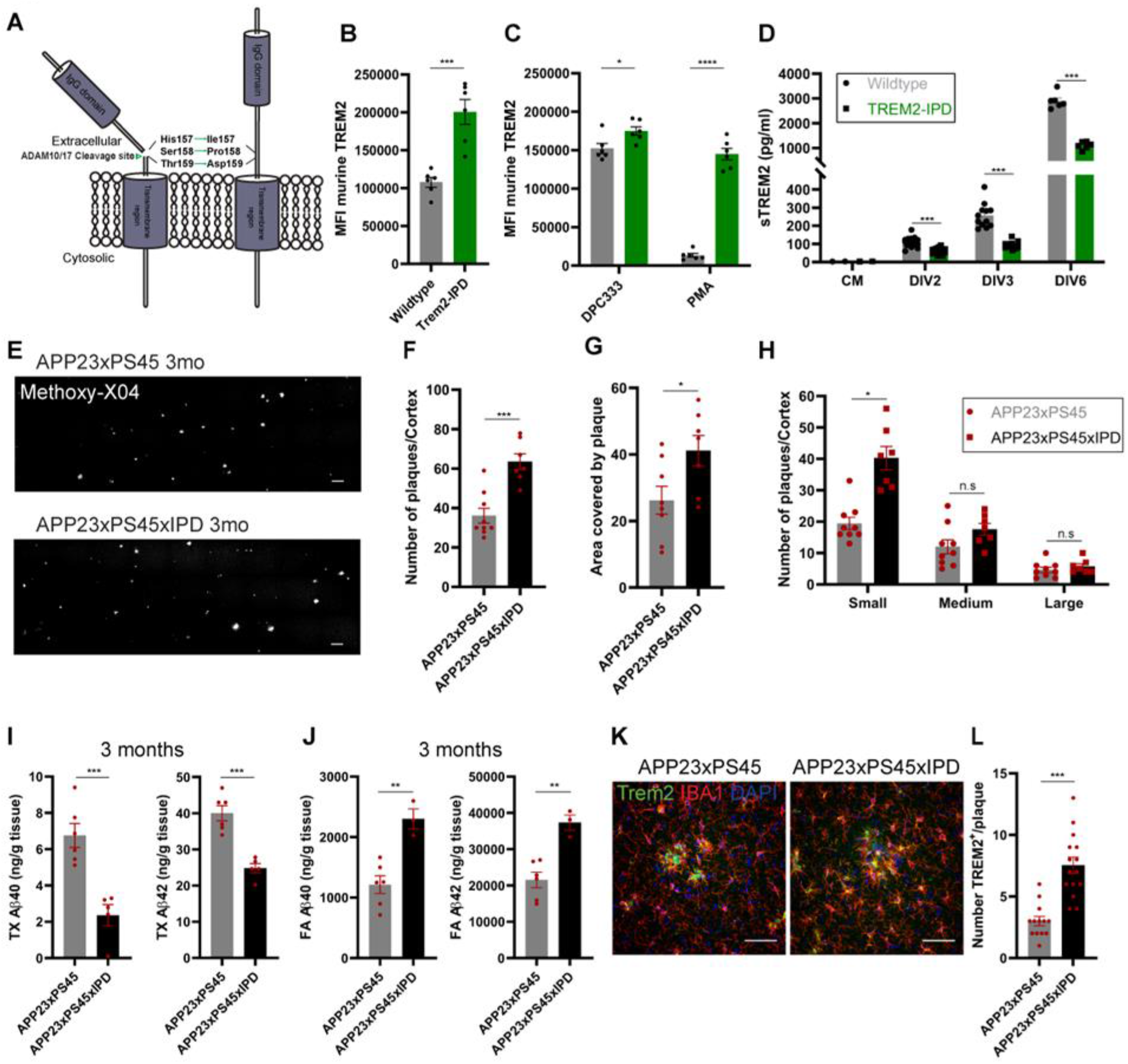
IPD substitution reduces membrane TREM2 cleavage and increases plaque deposition in a mouse model of AD. (A) Schematic showing the modification of amino acids on the stalk region of TREM2 from native HST to IPD to reduce proteolytic cleavage of TREM2. (B) Flow cytometric analysis of murine TREM2 on BMDM from WT and Trem2-IPD mice treated with DMSO (unpaired t-test; p=0.0004). (C) ADAM inhibitor DPC333 (unpaired t-test; p=0.0240) and PMA (unpaired t-test; p<0.0001, symbols represent technical replicates). (D) Levels of soluble TREM2 in the media collected on days in vitro 2, 3 and 6 from acute brain slice cultures of WT and Trem2-IPD mice as measured by ELISA (unpaired t-test; p<0.0001). (E) Representative images of Methoxy-X04 staining showing amyloid plaque load in APP23xPS45 (upper image) and APP23xPS45xIPD (lower image) mice at 3 months of age (3mo). (F) Quantification of total plaque numbers (unpaired t-test; p=0.0002), (G) Area covered by plaques (unpaired t-test; p=0.03) and (H) Plaque size (unpaired t-test; small: p=0.0001, medium: p=0.0860, large: p=0.3975) in the cortex upon Methoxy-X04 staining. (I) ELISA analysis of soluble (TritonX-100 fraction) Aβ40 and Aβ42 levels in forebrain lysates at 3mo (unpaired t-test; p=0.0009 and 0.0002). (J) ELISA analysis of insoluble (Formic Acid fraction) Aβ40 and Aβ42 levels in forebrain lysates at 3mo (unpaired t-test; p=0.0026 and 0.0023 respectively). (K) Representative images of TREM2 and IBA1 co-staining. (L) Quantification of images shows increased numbers of TREM2^+^ cell bodies around each plaque (unpaired t-test p<0.001) in the APP23xPS45xIPD mice. Scale bar for E=100μm and K=50μm. Error bars represent SEM.

### IPD-substitution increases Aβ plaque deposition at early stages in a mouse model of AD

To investigate the effect of IPD-substitution in the progression of AD, we crossed Trem2-IPD mice to an APP23xPS45 mouse model (Busche et al., 2008) to generate APP23xPS45xIPD mice. Analysis of Aβ plaque load in 3 months old (3mo) mice, age at which visible deposition of plaques begin revealed an increase in total plaque numbers (Figure 1E and 1F) and area covered by plaques (Figure 1G) in the cortex of APP23xPS45xIPD compared to APP23xPS45. In addition, quantification of Aβ plaque sizes shows this difference to stem from an increase in the number of small plaques indicating the appearance of newly formed plaques rather than plaque expansion (Figure 1H). We next determined the levels of soluble and insoluble Aβ40 and Aβ42 in 3mo mice. Compared to forebrain extracts of APP23xPS45 mice, APP23xPS45xIPD mice showed lower levels of soluble Aβ in the TritonX-100 (TX) fraction (Figure 1I). In contrast, levels of deposited insoluble Aβ in the formic acid (FA) fraction were higher in the APP23xPS45xIPD mice compared to APP23xPS45 (Figure 1J). However, quantification of Aβ plaques in the cortex of 7 months old (7mo) mice, when Aβ pathology is at the peak, exhibited no difference between the two genotypes in plaque numbers or size (Figure S2A-S2C). Similarly, while we noticed significant increase in TX soluble Aβ42, no differences were observed in insoluble Aβ40 and Aβ42 (Figure S2D) suggesting that plaque load plateaus by 7mo.

Impaired Trem2 function has been shown to affect microglial barrier formation around Aβ plaques and plaque compaction resulting in more toxic fibrillary plaques (Yuan et al., 2016a, Wang et al., 2016). To test if the increased Aβ load in 3mo APP23xPS45xIPD mice is reflected by plaque type, we stained sagittal brain sections with AbetaOC, a marker of fibrillary plaques. Analysis of AbetaOC staining over the Methoxy-X04 label showed no differences in area covered or in the volume of fibrillary plaques between APP23xPS45 and App23xPS45xIPD mice (Figure S3A). We assessed plaque associated microglia through immunofluorescence on the cortex of 3mo AD mice for TREM2 and IBA1 and observed more TREM2 positive microglia around the plaques (Figure 1K and 1L) and increased IBA1 volume (Figure S3B) in the APP23xPS45xIPD mice compared to the APP23xPS45. Moreover, using CD68 as a marker of activated microglia we found increased CD68 co-localization with Aβ plaques (Figure S3C) in APP23xPS45xIPD mice. This suggests that IPD-substitution induces increased microgliosis that facilitates plaque deposition at early stages of amyloid pathology in line with a recent study implicating DAM markers Axl and Mertk in plaque formation (Huang et al., 2021).

### Single cell RNA sequencing of amyloidosis mouse model identifies major cell types

To characterize the effect of TREM2 modification on microglia and other cell types in-vivo we employed single cell RNA sequencing (scRNAseq) of the cortex of WT, Trem2-IPD, APP23xPS45 and APP23xPS45xIPD mice at the beginning (3mo) and at peak (7mo) of Aβ pathology (Figure 2A). A total of 69,144 cells from 24 samples were embedded onto an Uniform Manifold Approximation and Projection (UMAP) plot to reveal nine major cell-types (Figure 2B). Clusters were annotated based on marker gene expression into microglia (Cl.1), oligodendrocytes (Cl.2), oligodendrocyte precursor cells (OPC; Cl.3), astrocytes (Cl.4), endothelial cells (Cl.5), choroid plexus (Cl.6), macrophages (Cl.7), vascular smooth muscle cells (Cl.8) and neurons (Cl.9) (Figure 2C). Remarkably, frequency analysis of each cluster across all samples at 3mo showed an increase in the abundance of microglial cluster in the Trem2-IPD mice under naive conditions and a further expansion with Aβ pathology. We also noticed a reduction in astrocyte abundance in AD mice while all other cell types were evenly represented (Figure 2D). At 7mo, we noticed similar representation of all cell types expect for a depletion of astrocytes and neurons in the AD mouse model due to the limitation of the dissociation method in capturing highly branched cell types.

**Figure 2:**
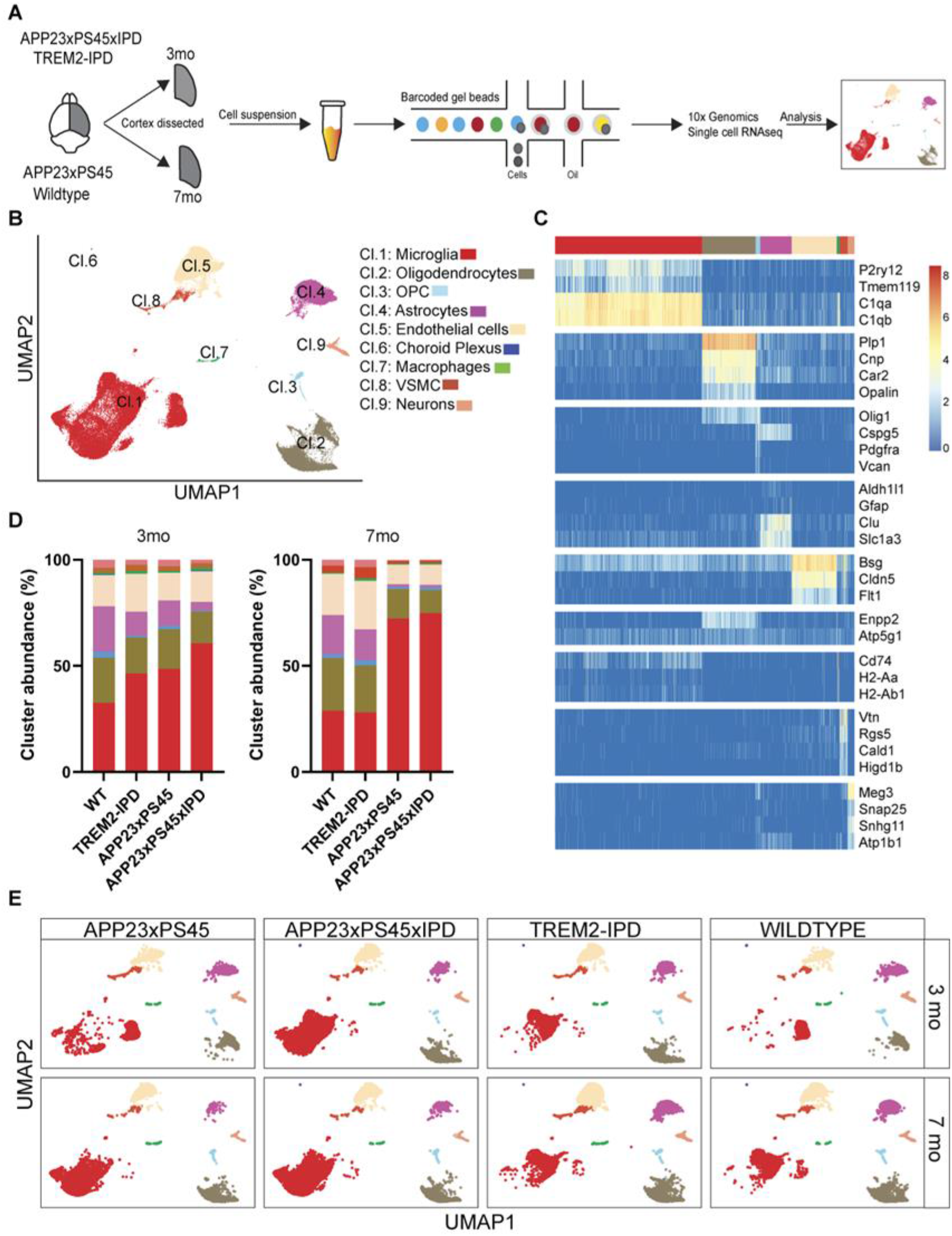
Single cell RNA sequencing of amyloidosis mouse model identifies major cell types. (A) Schematic diagram representing the workflow of scRNA-seq from the cortex of 3mo and 7mo wildtype and transgenic mice. (B) Uniform Manifold Approximation and Projection (UMAP) plot of all cells sequenced identifies 9 major clusters as annotated based on marker expression (24 samples, 69,144 cells derived from 3 independent mouse cortex per genotype and time-point as mentioned in (A). (C) Heat map showing the expression of markers used in annotating each cluster across all samples. (D) Stacked bar plots showing cell percentage composition of each cluster at 3mo and 7mo time points. (E) UMAP plots split by genotype per time point. All panels are derived from 3 mice per genotype and two time points.

### IPD-substitution shifts 3mo microglial clustering to resemble a late-stage state

To dissect IPD mediated microglial responses, we sub-clustered all microglia based on previously published datasets (Keren-Shaul et al., 2017, Sala Frigerio et al., 2019, Hammond et al., 2019, Krasemann et al., 2017, Masuda et al., 2020) into homeostatic (Cl. 1-2), DAM (Cl.3-4), Inflammatory DAM (Cl.5-6), Interferon response microglia (Cl.7), Spp1^+ve^ (Cl.8) and proliferating microglia (Cl.10) (Figure 3A and 3B). In addition to this, we also identified a unique cluster that resembled DAM but displayed downregulation of long non-coding RNA genes such as Malat1 and Xist (Cl.9), which we labelled as the Malat1^−ve^ cluster. Focusing on a per-cluster analysis, we discovered IPD-substitution to drive an age related shift in clustering. First, microglia of WT mice was predominantly homeostatic in nature expressing genes such as Tmem119, P2ry12, C1qa and were represented within Cl.1 (99.03%) and Cl.2 (94.91%) separated by age (early for 3mo and late for 7mo, respectively). Microglia from 3mo Trem2-IPD mice albeit transcriptionally similar to WT were largely represented within Cl.2 together with the 7mo mice (0.14% in Cl.1 and 98.52% in Cl.2; Figure 3C, 3D and S4A). No major differences were observed between 3mo and 7mo Trem2-IPD mice suggesting that IPD-substitution induces a mature microglial phenotype. Compared to WT mice, 3mo APP23xPS45 microglia showed a reduction in the fraction of cells within homeostatic cluster (63.51% in Cl.1) followed by enrichment in Cl.3 (7.21%) expressing DAM marker genes such as Trem2, Cst7, Cd68, Cd9, Tyrobp and Cl.5 (24.98%) expressing genes involved in immune regulation such as Ccl3, Ccl4, Il1β and MHC-II molecules like Cd74 and H2-D1. Intriguingly, microglia from 3mo APP23xPS45xIPD segregated separately into Cl.2 (62.92%), Cl.4 (3.07%) and Cl.6 (26.98%) very closely resembling microglia of 7mo mice.

**Figure 3:**
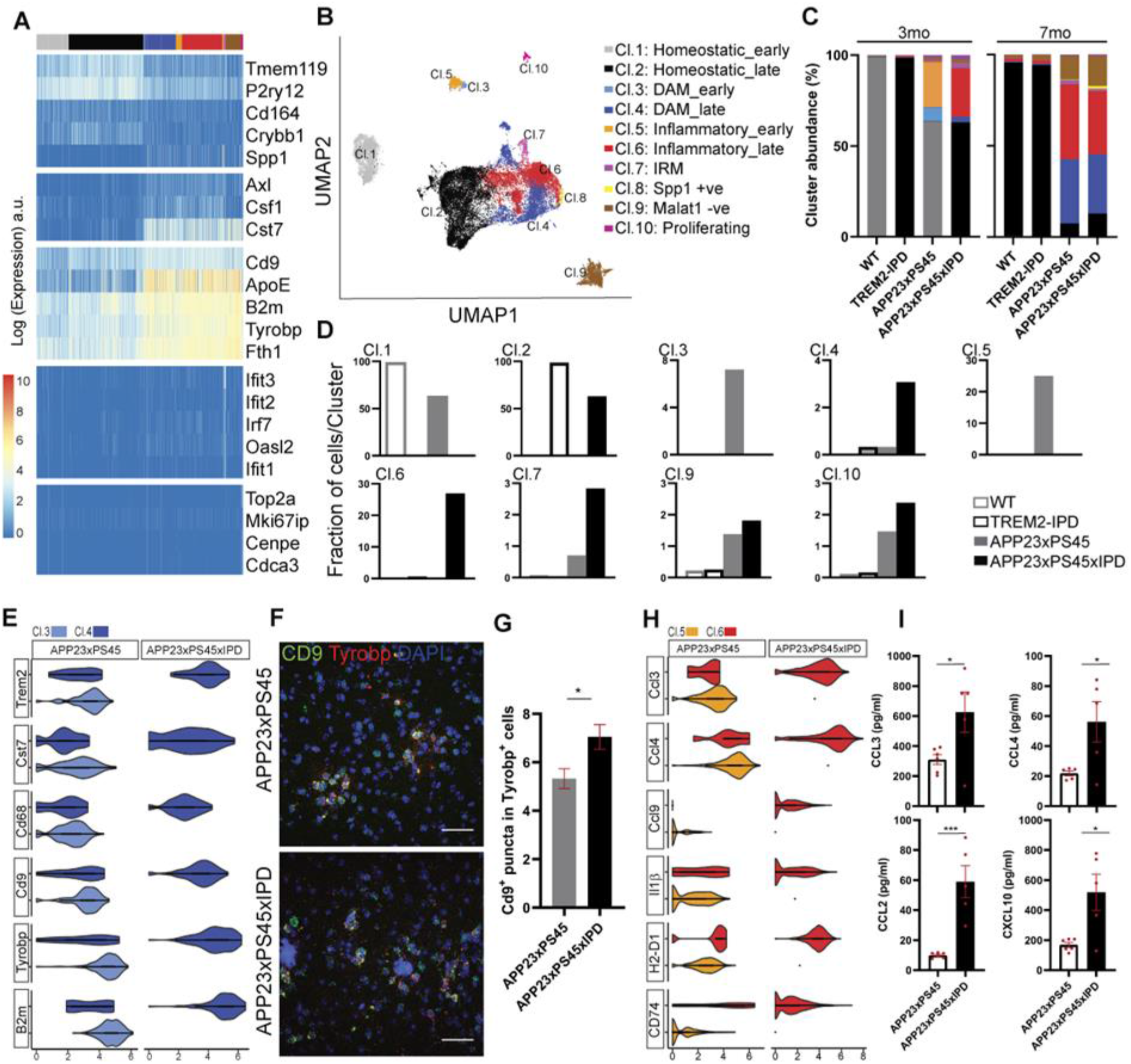
IPD-substitution shifts 3mo microglial clustering to resemble a late-stage state. (A) Heat map representation of genes used in sub-clustering of microglia from all samples. (B) UMAP visualization of microglial sub-clusters and the annotation of each cluster based on marker expression; early and late signifies clusters belonging to 3mo and 7mo mice respectively (n= 13,222 cells at 3mo and 19,305 cells at 7mo, obtained from 3 independent mouse cortex per genotype and time-point). (C) Stacked bar plots showing the percentage abundance of each microglial cluster across all four genotypes at 3mo and 7mo. (D) Fraction of cells belonging to each microglial cluster as labelled, displayed as bar plots from all samples at 3mo. Clusters 2, 4, 6 and 10 are composed of microglia predominantly from the IPD mice. (E) Violin plot representation of select DAM markers within Cl.3 and Cl.4 at 3mo; shape represents cell density and inserted bars represents median range. (F) Representative images co-stained for Trem2 and Iba1 (top panels) and RNAscope-ISH marked with probes for Cd9 and Tyrobp (bottom panels) highlighting distribution of DAM markers in APP23xPS45 and APP23xPS45xIPD mice. (G) Quantification of images shows increase in number of Cd9^+^ puncta within Tyrobp^+^ cells in APP23xPS45xIPD mice at 3mo (unpaired t-test p=0.010). (H) Violin plots of highly expressed genes related to cytokines, chemokines and MHC-related activation in Cl.5 and Cl.6. (I) quantification of cytokines and chemokines extracted from forebrain lysates of APP23xPS45 and APP23xPS45xIPD mice at 3mo. Unpaired t-test *p=0.01, ***p=0.007. Error bars represent SEM. Scale bar=50μm.

Alongside the DAM and inflammatory clusters we also observed the appearance of an interferon response microglia cluster (IRM; 2.83% in Cl.7) as characterized by the expression of Ifit2, Ifit3, Oasl2 and Irf7 (Figure 3C, 3D and S4A) that was absent in the APP23xPS45 mice (0.7%). Comparison of gene expression between Cl.3 and Cl.4 cells revealed elevated expression of a subset of DAM genes in the 3mo APP23xPS45xIPD mice compared to APP23xPS45 (Figure 3E). We next performed RNAscope-ISH for Cd9 and Tyrobp probes (Figure 3F) and found increased Cd9 puncta within Tyrobp^+^ cells (Figure 3G). Similarly, comparison of Cl.5 and Cl.6 revealed increase in transcripts encoding cytokines and trophic factors in 3mo APP23xPS45xIPD mice compared to APP23xPS45 (Figure 3H). In line with this, biochemical analysis on forebrain lysates also showed an increase in the levels of cytokines and chemokines in the APP23xPS45xIPD mice (Figure 3I). However, per-cluster analysis of microglia in 7mo mice showed minimal differences between genotypes with the exception of the appearance of Cl.8 expressing markers of inflammation such as Spp1 and Gpnmb in the APP23xPS45xIPD mice (Figure S5A). Furthermore, we also noticed a small expansion of the Malat1^−ve^ cluster (Cl.9) predominantly made up of ribosomal RPL, RPS family genes and mitochondrial (mt-Atp6, mt-Co2, mt-Nd1) genes in the APP23xPS45xIPD mice whose function still remains unknown. Thus, IPD-substitution drives microglia to a pronounced activated state in 3mo APP23xPS45xIPD mice resulting in upregulation of inflammatory genes compared to the APP23xPS45 mice.

### Pseudotime analysis reveals separate fates induced by IPD-substitution

To address the differences in microglial clustering between the WT and IPD mice, we mapped the trajectory of microglia at 3mo from all four genotypes along a pseudotime axis. With the root set to homeostatic microglia of WT mice, cells were partitioned into 7 states (Figure 4A and 4B). Interestingly, the presence of IPD bifurcated the branch with microglia of Trem2-IPD and APP23xPS45xIPD following a separate trajectory with a terminal state at t7 (t=11[a.u]) compared to WT and APP23xPS45 mice (t5=6.1[a.u]) (Figure 4C). Distribution of microglial sub-clusters along the trajectory showed cells from Cl.1 were distributed between states t1, t3 and t5 while Cl.3 and Cl.5 were present within t5. Cells of Cl.2 instead were distributed between t4, t5 and t7 which was also composed of cells from Cl.4, Cl.6 and Cl.7 (Figure 4D and 4E). Moreover, we assessed the longitudinal progression of microglia into DAM or IRM states from 3mo and 7mo in AD mice. Using B2m, Cd9, Fth1 and Trem2 as markers of DAM and Ifit2, Ifit3, Irf7 and Oasl2 as cluster specific markers of IRM we found that IPD-substitution accelerates microglial transition to DAM (Figure 4F) and IRM states (Figure 4G). Taken together, these results suggest that IPD-substitution primes microglia to a mature phenotype under naive state that can then transform more rapidly into heterogeneous activated states under pathological conditions compared to native-TREM2.

**Figure 4:**
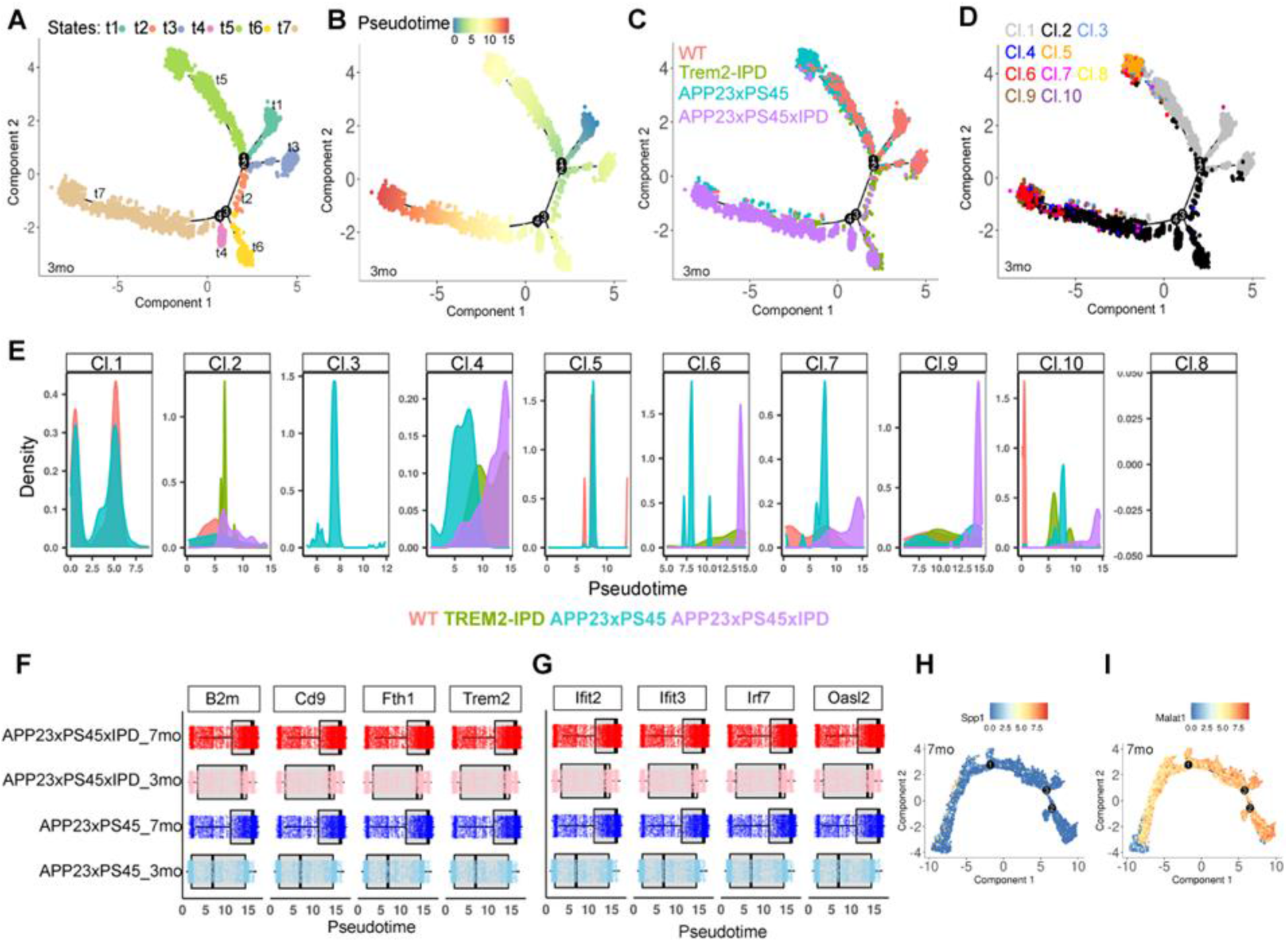
Pseudotime analysis reveals separate fates induced by IPD-substitution. Cell trajectory analysis of microglial sub-clusters using Monocle at the 3mo time-point based on (A) States (t1=0.6, t2=3.7, t3=5.1, t4=6.1, t5=6.1, t6=6.7, t7=11). (B) Pseudotime (with root set to homeostatic microglia in 3mo WT mice). (C) Genotype and (D) Sub-clusters (as established in Fig.3b) showing the bifurcation of microglial progression into two separate axis driven by IPD-substitution. (E) Density distribution plot of the pseudotime showing microglia sub-clusters across all genotypes at 3mo. (F) Bar plots showing pseudotime progression of microglia from homeostatic to DAM state in APP23xPS45 and APP23xPS45xIPD mice at 3mo and 7mo. (G) Bar plots showing pseudotime progression of microglia from homeostatic to IRM state in APP23xPS45 and APP23xPS45xIPD mice at 3mo and 7mo. (H) Expression of SPP1 plotted over the trajectory tree in 7mo mice. (I) Expression of Malat1 plotted over the trajectory tree in 7mo mice.

In addition, pseudotime analysis of 7mo across all genotypes revealed that WT and Trem2-IPD mice progressed from a homeostatic state to an activated state along the same trajectory (Figure S5B-S5F). In accordance with Figure 3C we found the appearance of SPP1^+ve^ cluster in the APP23xPS45xIPD mice with increasing expression towards the end of pseudotime axis (Figure 4H) and a small expansion of Malat1^−ve^ cluster (Figure 4I) at the farthest end of the pseudotime.

### Reactive oligodendrocytes appear in AD mice and are unaffected by IPD-substitution

We expanded our analysis to other cell types. On the one hand, we noticed an age related shift in basal states of astrocytes and oligodendrocytes in non-AD mice similar to microglia (Figure 2E) possibly suggesting an effect of microglial activation in the maturation of other glial cells. On the other hand, only oligodendrocytes underwent the most notable transformation during the course of Aβ pathology with the appearance of a ‘reactive’ cluster (Zhou et al., 2020) (Cl.5) with upregulation of genes such as C4b, Serpina3n, B2m, Il33, Ifi27 (Figure S6A-S6C) in 7mo AD mice. Comparison of gene expression across all genotypes using C4b, Serpina3n and B2m as Cl.5 specific markers yielded an increase in levels of the three genes upon Aβ pathology whereas IPD-substitution had no effect on this phenotype (Figure S6D). We corroborated this finding using RNAscope-ISH for Serpina3n and immunofluorescence for C4b that showed that majority of the expression lie within oligodendrocytes while no differences in expression were observed between the APP23xPS45 and APP23xPS45xIPD mice (Figure S6E-S6F). The ‘reactive oligodendrocytes’ were however absent in 3mo mice (Figure S6G).

### IPD-substitution induces neurite dystrophy at 3mo

Finally, to determine the effect of the microglial response on neuronal health, we performed immunofluorescence for APP as a marker of dystrophic neurites on 3mo and 7mo AD brain tissues. Compared to 3mo APP23xPS45 mice, APP23xPS45xIPD mice exhibited higher number of APP^+^ puncta around the plaques (Figure S7A-S7B). While we observed an age related increase in APP^+^ staining around each plaque at 7mo, no differences between the two genotypes were observable (Figure S7C-S7D). Similarly at 3mo and 7mo we found no differences in the volume of APP^+^ staining.

## Discussion

Since GWAS studies identified variants of TREM2 in AD, there has been urgency in understanding TREM2 biology and developing therapeutic strategies. In rodents, loss of function (LOF) models such as Trem2^-/-^and Trem2^R47H^ mice suggested a therapeutic hypothesis aimed at increasing microglial TREM2 function. In this study, we generated a transgenic mouse model harboring amino acid substitution (Trem2-IPD) of the ADAM10/17 protease recognition site on the stalk region of TREM2 in an attempt to reduce TREM2 shedding and to increase TREM2 function in microglia. In doing so, we investigated how sustained reduction of TREM2 shedding and increased membrane load affected myeloid cell responses. Using a double transgenic mouse model of AD in which rapid deposition of Aβ plaques appear from 3 months of age onward, we show for the first time that increasing TREM2 stabilization facilitates Aβ deposition and neuroinflammation at early stages of the pathology.

Increased neuroinflammation resulting from microglial and astrocytic activation has been postulated to initiate and to accelerate Aβ deposition in several APP transgenic mouse models (Qiao et al., 2001, Holtzman et al., 2000, Minter et al., 2016). Here, we employed scRNAseq to investigate how IPD-substitution affects the cellular landscape of neuroinflammation. We demonstrated that microglia transform from a homeostatic or resting state to a heterogeneous activated state from 3mo to 7mo expressing transcripts related to inflammation, phagocytosis, cytokine or chemokine expression, interferon signature and a Malat1^−ve^ population of yet unknown function. TREM2 modification in this AD mouse model accelerated the microglial shift into four different activated cell states (DAM, Inflammatory DAM, IRM and SPP1^+ve^). We corroborated these findings by performing cytokine analysis on forebrain lysates revealing an increase in inflammation within the brain of transgenic mice.

Compared to studies showing that TREM2 ablation or LOF affects microglial maturation, locking microglia in a homeostatic state preventing them from responding to extracellular signals (Mazaheri et al., 2017, Keren-Shaul et al., 2017, Yuan et al., 2016b), our data suggests that increased TREM2 function accelerates microglial response to Aβ pathology. Interestingly, we observed minimal changes in microglial gene signature between 3mo and 7mo upon IPD-substitution in AD suggesting that microglia activation through TREM2 stabilization reaches a saturation point.

Moreover, our data shows that reducing TREM2 shedding triggers an upregulation of inflammatory genes and aggravates plaque load and APP^+^ dystrophic neurites at early stages of the disease. One possible explanation for this could be that constitutive reduction of TREM2 shedding primes microglia earlier on during development that persists into adulthood. This is in line with our observation that microglia exhibit a mature phenotype wherein 3mo Trem2-IPD microglia cluster with 7mo WT mice with no significant differences in gene signature. Indeed, it has been shown that lack of TREM2 activation during development impairs microglia maturation and results in the development of social deficits and altered synaptic connectivity in adult mice (Filipello et al., 2018) and PLOSL in humans (Bianchin et al., 2004). Furthermore, in light of the therapeutic modalities based on TREM2 activating antibodies (Wang et al., 2020, Schlepckow et al., 2020), our study provides insights into the effects of chronic or early intervention placing an importance on the timing of treatment in AD.

Alongside effects on microglia, we also observe an age related shift in clustering of astrocytes and oligodendrocytes. Furthermore, we also confirm the presence of reactive oligodendrocytes characterized by the presence of markers such as Serpina3n and C4b in the cortex of 7mo double transgenic mice. Interestingly, IPD-substitution had no effect on the appearance of reactive oligodendrocytes suggesting that they are independent of TREM2 activity. Future studies on the role of Trem2-IPD in development and ageing will further advance our understanding of the effects caused by chronic TREM2 stabilization.

## Acknowledgements

We thank Sven Schuierer for help in analyzing the sequencing data. We thank Daniel Breustedt, Hong Lei, Thomas Nicholson and Wibke Schwarzer for help in the generation of the Trem2-IPD mouse model. We thank Noemie Siccardi for the technical support of the RNAscope-ISH experiments.

## Author contributions

R.D and M.N designed experiments, analyzed and interpreted the data and co-wrote the manuscript. M.B. and I.B. designed and performed the brain slice experiments and analyzed data. I.B. and U.N. designed and performed biochemical experiments and analyzed data. T.S., S.R., D.R.S. and F.G. designed and performed cytokines and chemokines measurements and analyzed data. S.J., R.B. and D.F. designed and performed BMDM experiments and analyzed data. R.D. and M.B. performed dissections and dissociations for the scRNAseq experiments and primary microglia experiments and analyzed data. R.D. designed and performed RNAscope-ISH experiments and staining experiments and analyzed data. A.W., R.C., U.N. and G.R. designed and performed sequencing experiments. M.N. and C.G.K designed and analyzed scRNAseq data. D.F., D.R.S., U.N. and F.G. designed and supported interpretation of the Trem2-IPD mouse model. I.G. conceived the study, designed experiments, interpreted the data and co-wrote the manuscript.

## Data and code availability

Single cell sequencing data related to this study is available at the NCBI Sequence Read Archive with the project ID PRJNA714499.

## Experimentl model

### Animals

APP23xPS45 double transgenic mice were generated as previously described (Busche et al., 2008) by crossing mice overexpressing human APP carrying Swedish double mutations at positions 670/671 (APP23) with mice harboring a G384A mutation in the human Presenilin1 gene (PS45). To generate the Trem2-IPD knock-in (KI) mice, two guide RNAs (gRNAs) and a donor oligo (200bp) containing the mutant amino acids IPD: ATCCCCGAC replacing the wildtype amino-acids HST: CACAGCACC were designed to target the mouse TREM2 allele. These gRNAs, #1 (complementary sequence: 5’-gggaccactactgtacctgg), #2 (forward sequence: 5’-aagtggaacacagcacctcc) were ordered as crRNAs from IDT (IA, USA) as part of their Alt-R CRISPR/Cas9 system. sgRNA1 or sgRNA2 was microinjected, along with the universal Alt-R tracrRNA (IDT, IA, USA), donor oligo and Cas9 protein (PNA Bio, CA, USA) into the pro-nucleus of fertilized C57Bl/6J oocytes. Viable 2-cell embryos were re-implanted into pseudo-pregnant B6CBAF1 females. The resulting pups were genotyped by PCR (primer F1: 5’-agctacccgctactgcaaag, primer R1: 5’-cccgatgagctcttccacat; expected product length: 635bp; PCR program: 95°C 3min, 95°C 20sec, 60°C 20sec, 72°C 45sec, for 35 cycles, 72°C 5min, 4°C forever), XhoI digested and Sanger sequenced. Eight KI founders were identified and back-crossed with C57BL/6J (JAX, 000664) and the F0 and F1 generation was subjected to 1) MiSeq (Illumina Inc, San Diego, USA) sequencing using forward primer 5’-atgctggagatctctgggtcc and reverse primer 5’-gtgagttgctacaaagggctcc to generate amplicons and 2) NuGEN’s Amplicon Sequencing System (part number 9092-256, Tecan, Redwood City, USA) for Next-Generation Sequencing library construction. Furthermore, two founder lines were chosen and bred to homozygosity. One homozygous of each founder line was then subjected to targeted locus amplification (TLA) at Cergentis (Utrecht, The Netherlands) to detect integration of the transgene in mouse chr17:48,351,169-48,351,193 as intended. All fusion reads at the integration site were similar and no wildtype reads were found at the integration site. Mice were genotyped using the following primers: Knock-in with primer sets 778-Trem2-CR-6 (5’-gatagggaatcgaccagaggc) and 774-Trem2-CR-2 (5’-actgtacct cgagtcggggat); wildtype primer sets 775-Trem2-CR-3 (5’-caagtggaacacagcacctcc) and 777-Trem2-CR-5 (5’-aggtatgtggctgctcag gg). Expected KI band size: 246bp, wildtype band size: 350bp. PCR program: 95°C 3min, 95°C 30sec, 64°C 20sec, 72°C 30sec, step 2 for 35 cycles, 72°C 5min, 8°C forever.

APP23xTrem2-IPD mice and PS45xTrem2-IPD mice were crossed to obtain APP23xPS45xIPD mice. Female mice (n=3 per genotype and time point) were used in all single cell RNA sequencing experiments. Both male and female mice were used for validation. All animals were housed in a temperature-controlled room and maintained on a 12hr light/dark cycle. Food and water were available *ad libitum* and experiments were conducted in accordance with the authorization guidelines for the care and use of laboratory animals of Basel City, Switzerland.

## Method details

### Culture of Bone marrow derived macrophages

Bone marrow derived macrophages (BMDM) were prepared from 3 adult WT and Trem2-IPD mice. Tibia and the femur bones were dissected clean of surrounding muscle tissue and the marrow were flushed out using a 30G needle fitted onto a 10ml syringe. Cell suspension was filtered through a 100μm cell strainer and incubated for 5 minutes in Red blood cell lysis buffer. Following centrifugation, BMDM were washed with PBS and resuspended in a RPMI1640 culture media containing 10% FBS (Gibco, 10082147), Pen/Strep (Gibco, 15070063), 1% Sodium pyruvate (Gibco, 11360-070), 1% NEAA (Gibco, 11140035), 0.025M HEPES (Gibco, 15630080), 50μM β-Mercaptoethanol and 40ng/ml M-CSF (R&D systems, 416-ML-050). Three days after seeding, cells were replenished with media containing 40ng/ml M-CSF, 20ng/ml M-IL4 (R&D systems, 404-ML-050) and 20ng/ml M-IL13 (R&D systems, 413-ML-005). BMDM were used in experiments after 1week of differentiation.

### FACS analysis of membrane TREM2

For measurement of cell surface TREM2 expression, BMDM were treated with either 5μm DPC333 (Qian et al., 2007) O/N or with 50ng/ml PMA (Sigma Aldrich, 19-144) for 30 minutes. Cells were detached using 1X TrypLE enzyme (ThermoFisher, 12605010) and incubated with FC block (anti-CD16/32; eBioscience, 14-0161-85) in FACS buffer (PBS containing 2% FBS, 0.5mM EDTA and 0.05% Sodium azide) for 20 minutes at 4°C following which cells were stained in suspension for anti-mouse TREM2 biotinylated (R&D systems, BAF1729) for 30 minutes at 4°C. Streptavidin-AF488 secondary antibody was added for 30 minutes at 4°C. Staining were analyzed in a FACS canto upon gating for LIVE/DEAD Fixable aqua (Molecular probes, L34966) to remove dead cells.

### Cell survival assay

Differentiated BMDM were plated in 96-well plate at density of 30,000 cells/well either in the presence of 40ng/ml M-CSF (R&D systems, 416-ML-050) or without M-CSF. After 10 days, cellular ATP levels were measured using CellTiter Glo (Promega, G755A).

### Acute slice preparation

6 week old female wild-type and TREM2-IPD mice (n=3) were decapitated and brains were dissected into freshly prepared dissection medium containing 2X MEM (Gibco, 04195120M), 25mM HEPES, 10mM Tris (hydroxymethyl) aminomethane and 1% Pen/Strep. Sagittal sections of 150μm thickness were cut using a vibratome (Leica, VT1000S) and transferred to insert wells (Millicell Cell Culture Insert, 30mm, Millipore) in a 6-well plate filled with culture medium containing 2X MEM (Gibco, 04195120M), 25% HBSS, 25mM HEPES, 10mM tris(hydroxymethyl) aminomethane, 25% Horse serum and 1% Pen/Strep. The entire procedure was done on ice with precooled solutions until culturing. Sections were incubated for 6 days in an incubator (35°C, 5% CO2). For sTREM2 measurements, 500μl of the culture medium was collected at day in-vitro 2, 3 and 6. 3 wells per mice per genotype was collected and ELISA for Trem2 was performed as mentioned below.

### Measurement of TREM2

For sTREM2 measurements, multi array 96-well plates (MSD, L15XA-3) were coated with mouse anti-TREM2 (R&D systems, AF1729) capture antibody O/N at 4°C. Plates were incubated with blocking buffer containing 2% BSA and 0.05% Tween20 in 1xTBS for 1 hour at RT and washed with 0.05% Tween20 in 1xTBS. Culture medium from acute brain slices were collected, spun down at 1000RPM for 5 minutes and 25μl of the supernatant was added onto the coated 96-well plates. Following incubation with the detection antibody (biotinylated anti-TREM2; R&D systems, BAF1729) for 1hour at RT and Streptavidin SULFO TAG™ (MSD, R32AD-1), TREM2 levels were quantified using a MSD Sector Imager 600.

### Primary microglial culture

Microglia were harvested from mixed-glial cultures prepared from Wildtype and Trem2-IPD newborn mice as described previously (Prinz et al., 1999) with some modifications. Brains from 9-10 pups at postnatal day 7 were dissected and brainstem, olfactory bulb, meninges removed. Tissue was chopped into small pieces and trypsinized in Hibernate-A media containing 1X Trypsin (Sigma, T4549) for 20 minutes at 37°C followed by mechanical dissociation using a Pasteur pipette in Hibernate-A media containing DNAse1 (Life Technologies, 18047019). Cells were filtered through a 70μm cell strainer and cultured in microglia medium containing DMEM supplemented with GlutaMAX (ThermoFisher, 31966-021), 10% FBS, 1X Pen/Strep, 1X Sodium Pyruvate (Gibco, 11360070) and 1x nonessential amino acids (Gibco, 11140050). Microglia were dislodged from the mixed-glial cultures 10-14 days later by vigorous shaking.

### Phagocytosis by primary microglia

For in-vitro phagocytosis, primary microglia from Wildtype and Trem2-IPD mice were plated in Poly-L-lysine coated 96-well plates at seeding density of 40,000 cells/well and let attach for 3 hours. pHrodo™ Green *S. aureus* Bioparticles (ThermoFisher, P35367; 3μg/ml), pHrodo™ Green *E.coli* Bioparticles (ThermoFisher, P35366; 3μg/ml), pHrodo™ Red LDL (ThermoFisher, L34356; 1μg/ml) and pHrodo™ Green Tau (produced at Novartis; 1μg/ml) were diluted in microglia medium and added to individual wells. The plates were placed in an IncuCyte S3 live cell analysis system and four images per well were acquired every hour for 9 hours. The integrated fluorescence intensities were extracted for each well and analyzed in a GraphPad Prism software.

### Biochemical characterization of Aβ

Human Aβ40 and Aβ42 levels were quantified using Human (6E10) Abeta Peptide Ultrasensitive Kits (Meso Scale Discovery, N451FSA-1). Forebrain homogenates of APP23xPS45 and APP23xPS45xIPD mice were prepared by sonication in a solution containing 10x w/v TBS and 1x Halt TM Protease & Phosphatase (PP) Inhibitor Cocktail (ThermoFisher, 78444). For the soluble fraction, 50μl of the homogenate was treated with 2% Triton-X and incubated for 15 minutes on ice. The mixture was centrifuged at 45,000RPM for 15 minutes. For the insoluble fraction, 50μl of the homogenate was treated with 70% Formic Acid for 15 minutes followed by neutralization in 1M Tris-base at pH 10.85 O/N at 4°C. The mixture was centrifuged at 14,000RPM for 15 minutes and the supernatant was used for Aβ measurements.

### Cytokine measurements

Mouse CCL2 (MCP-2), CCL3 (MIP1a), CCL4 (MIP1b) and CXCL10 (IP-10) were measured using a MSD-96-well Uplex 7-assay (Meso Scale Discovery, N05231A-1). Briefly, forebrain homogenates were treated with 10x RIPA lysis buffer (Millipore, 20-188) and 1x Halt TM Protease & Phosphatase (PP) Inhibitor Cocktail (ThermoFisher, 78444) for 10 minutes at 4°C. The mixture was centrifuged at 18,000RPM for 10 minutes and the supernatant was collected for analysis.

### Cortical dissociation for scRNAseq

Mice were anesthetized under 3.5% Isoflurane and upon deep narcosis, transcardially perfused with ice-cold PBS for 3 minutes. The brains were harvested and the cortex was carefully dissected from a single hemisphere and placed in cold Hibernate-A solution (Life Technologies, A1247501). One cortical hemisphere from every mice was dissociated into single cells using an Adult Brain Dissociation kit (Miltenyi Biotech, 130-107-677) following manufacturer’s instructions. In short, cortex was chopped into fine pieces and mechanically digested in GentleMACS C-tubes containing enzyme mixes 1 and 2. Cell clusters were removed by filtering through a 70um cell strainer followed by debris removal and red-blood cell lysis. No further myelin removal was performed. Dissociated cells were diluted in ice cold PBS containing 0.04% BSA with a density of 1000-1200 cells/μl.

### Single cell RNA-sequencing

Cellular suspensions were loaded on a 10x Genomics Chromium Single Cell instrument to generate single cell GEMs. Single cell RNA-Seq libraries were prepared using GemCode Single Cell 3’ Gel Bead and Library Kit according to CG00052_SingleCell3'ReagentKitv2UserGuide_RevD. GEM-RT was performed in a Bio-Rad PTC-200 Thermal Cycler with semi-skirted 96-Well Plate (Eppendorf, P/N 0030 128.605) following: 53°C for 45min, 85°C for 5min; held at 4°C. After RT, GEMs were broken and the single strand cDNA was cleaned up using DynaBeads^®^ MyOne™ Silane Beads (Life Technologies, P/N 37002D). cDNA was amplified using a Bio-Rad PTC-200 Thermal cycler with 0.2ml 8-strip non-Flex PCR tubes, with flat Caps (STARLAB, P/N I1402-3700): 98°C for 3min; cycled 10 to 12x: 98°C for 15s, 67°C for 20s, and 72°C for 1min; 72°C for 1min; held at 4°C. Amplified cDNA product was cleaned up with the SPRIselect Reagent Kit (0.6X SPRI). Indexed sequencing libraries were constructed using the reagents in the Chromium Single Cell 3’ library kit V2 (10x Genomics P/N-120237), following: 1) Fragmentation, End Repair and A-Tailing; 2) Post Fragmentation, End Repair & A-Tailing Double Sided Size Selection with SPRIselect Reagent Kit (0.6X SPRI and 0.8X SPRI); 3) adaptor ligation; 4) post-ligation clean up with SPRIselect (0.8X SPRI); 5) sample index PCR using the Chromium Multiplex kit (10x Genomics P/N-120262); 6) Post Sample Index Double Sided Size Selection- with SPRIselect Reagent Kit (0.6X SPRI and 0.8X SPRI). The barcode sequencing libraries were quantified using a Qubit 2.0 with a Qubit™ dsDNA HS Assay Kit (Invitrogen P/N Q32854) and the quality of the libraries were performed on a 2100 Bioanalyzer from Agilent using an Agilent High Sensitivity DNA kit (Agilent P/N 5067-4626). Sequencing libraries were loaded at 10 to 12pM on an Illumina HiSeq2500 with 2 × 50 paired-end kits using the following read length: 26 cycles Read1, 8 cycles i7 Index and 98 cycles Read2. The CellRanger suite (Zheng et al., 2017) was used to generate the aggregated gene expression matrix.

### Analysis of scRNA-seq

The single cell analysis was performed following a modular workflow assembled in R version 4.0.3 (Wegmann et al., 2019). The UMI count matrix from the 4 genotypes and 2 time points (n=3, 24 samples) included a total of 87,672 sequenced cells. Cells with UMI count lower than 1024 and number of detected genes less than 512 were filtered out. In addition, cells with percentage of UMIs mapping to higher than 20% of mitochondrial genes were excluded. Lowly expressed genes with fewer than 2 UMIs in less than 2 cells were also filtered out. The UMI count matrix after filtering included a total of 69,144 cells and 15,611 Ensembl gene IDs.

The data were normalized with the *scran* R package version 1.18.3 by deconvoluting size factors from cell pools with a size of 20, 100 and 5 cells and log2 transformed (Lun et al., 2016). Dimension reduction was further applied to reduce the technical noise: after subset the matrix with the 2,192 highly variable genes identified by the distance-to-median approach (Kolodziejczyk et al., 2015) ‘modelGeneVar’ and ‘getTopHVGs’ functions from *scran*) a principal component analysis was applied and the first 27 principal components accounting for 30% of the total variation across the cells were retained.

2D projection for cell visualization (Figure 2) was computed using the Uniform Manifold Approximation and Projection, UMAP (‘umap’ version 0.2.7.0, R package), using the Eulidean metric, umap-learn method(McInnes et al., 2018) version 0.4.6, python 3.7 and the size of the local neighborhood set to 100 (n_neighbors).

A graph-based clustering method from the *scran* R library was used to generate cell connection based on the nearest neighbors method (n_ neighbors=100). Community structure via Louvain clustering (Blondel et al., 2008) (*igraph* R package version 1.2.6) was further applied to find densely connected sub-graphs.

The signatures were assigned based on known cell population markers, as Microglia (P2ry12, Tmem119, C1qa, C1qb), Oligodendrocytes (Plp1, Cnp, Car2, Opalin), OPC (Olig1, Cspg5, Pdgfra, Vcan), Astrocytes (Aldh1l1, Gfap, Clu, Slc1a3), Endothelial cells (Bsg, Cldn5, Flt1), Choroid Plexus (Ttr, Enpp2, Atp5g1), Macrophages (Cd74, H2-Aa, H2-Ab1), VSMC (Vtn, Rgs5, Cald1, Higd1b) and Neurons (Meg3, Snap25, Snhg11, Atp1b1).

### Analysis of microglia and oligodendrocytes

The cells with microglia and oligodendrocytes signatures as previously assigned were analyzed individually. The UMIs were processed using the same criteria explained in the above paragraph, *i.e.* after filtering and cell normalization, a dimension reduction was performed for microglia (2,796 highly variable genes(Kolodziejczyk et al., 2015) further reduced by the first 54 principal components accounting for 10% of the total variation across the cells) and for oligodendrocytes (1,624 highly variable genes further reduced by the first 31 principal components accounting for 18% of the total variation across the cells).

The 2D UMAP projections (McInnes et al., 2018) on the reduced matrices were applied using parameters n_neighbors=10 for both microglia and oligodendrocytes dataset. The signatures were assigned based on known cell population markers: Microglia: Homeostatic (Tmem119, P2ry12, Cd164, Crybb1), DAM (Spp1, Axl, Csf1, Cst7), Inflammatory (Cd9, ApoE, B2m, Tyrobp, Fth1) IRM (Ifit3, Ifit2, Irf7, Oasl2, Ifit1) and Proliferating (Top2a, Mki67ip, Cenpe, Cdca3). Oligodendrocytes: OPC (Olig1, Cspg5, Pdgfra, Vcan), mature oligodendrocytes (Plp1, Cnp, Car2, Opalin) and Reactive oligodendrocytes (C4b, Serpina3n, B2m, Il33).

The Microglia trajectory analysis at 3mo and 7mo were computed using monocle (Trapnell et al., 2014) version 2.18.0 R package, using as input the counts of the top 2,796 high variable genes. The differential gene expression analysis was performed with Limma-trend (Ritchie et al., 2015), limma version 3.46.0 R package.

### Amyloid plaque staining and quantification

Paraffin embedded brain tissues were sectioned at 3μm thickness, deparaffinized in Xylene and dehydrated through a series of ethanol treatment steps. Following antigen retrieval in Citrate buffer (pH6; Quartett, 400300692), slides were incubated with blocking solution containing 5% serum (Goat or Donkey) and 2% BSA in 0.1% Tween-20 in PBS for 1 hour and subsequently with anti-amyloid fibrils OC antibody in blocking solution overnight at 4°C. Alexa-fluorophore conjugated secondary antibodies were added in blocking solution for 1 hour at 4°C followed by treatment with Methoxy-X04 (Tocris, 4920) for 15 minutes at room temperature to label amyloid-beta plaques. Confocal tiles of the entire cortical region were acquired with a LSM700 microscope and plaque numbers, volume, area covered and diameter were calculated using the ‘Surfaces’ function of Imaris. For plaque size analysis, objects <20μm in diameter were classified as small, 20-50μm as medium and >50μm as large.

### Immunofluorescence

Sagittal brain sections were permeabilized in 0.3% Triton-X in PBS for 20 minutes and incubated with blocking solution containing 5% serum (Goat or Gonkey) and 2% BSA in 0.1% Tween-20 in PBS for 1 hour and subsequently with primary antibodies in blocking solution overnight at 4°C. Alexa-fluorophore conjugated secondary antibodies were added in blocking solution for 1 hour at 4°C. Sections were then labelled for Methoxy-X04 (10 minutes at RT) or counterstained with DAPI and mounted with prolong gold anti-fade medium (Invitrogen, P36930). Images were acquired using a Zeiss LSM700 confocal microscope with the Zen 2010 Software package. For IBA1, CD68 and Trem2 staining: whole brain was dissected from PBS-perfused mice, each hemisphere separated and fixed with 4% PFA overnight at 4°C. 40μm thick sagittal sections were cut using a Vibratome (Leica, VT1200s) and processed as free floating sections. For C4b and Olig2 staining: each hemisphere of PBS-perfused mouse brains was placed in cryo-molds and embeded in Tissue Tek O.C.T compound. 10-12μm sagittal sections were cut onto Superfrost^+^ slides using a Cryostat (Leica, CM3050S).

Cell body volume of IBA1 was measured using the ‘Surpass’ function of Imaris. For CD68^+^/MethoxyX04^+^ quantifications, ‘Coloc’ function of Imaris was used. For dystrophic neurite quantifications, 3μm thick FFPE brain sections were deparaffinized in Xylene and antigen retrieval performed using citrate buffer. Staining for mouse anti-APP were carried out as mentioned above. Tiff images of individual channels were quantified using a Cellprofiler V2.2.0 software by identifying objects 12-30 pixels in diameter around each MethoxyX04 labelled plaque. Total number of APP^+^ neurites and the volume of neurites were calculated using the MeasureObjectSizeShape module.

List of primary antibodies used: Rabbit anti-amyloid fibrils OC (1:1000; Merck AB2286), Rabbit anti-Iba1 (1:1000; Wako, 019-19741), Sheep anti-Trem2 (1:500; R&D systems, AF1729), Rat anti-CD68 (1:500; Biorad, MCA1957), Goat anti-Olig2 (1:300; R&D systems, AF2418), Rat anti-C4b (1:100; Hycult, HM1046), Mouse anti-APP (1:100; Sigma, MAB348). Alexa-fluorophore conjugated secondary antibodies were used.

### RNAscope in-situ hybridization

Fluorescent in-situ hybridization was performed on OCT embedded fresh-frozen brain tissues from adult mice using RNAscope^®^ Fluorescent Multiplex Manual Assay Kit (Advanced Cell Diagnostics) following manufacturer’s instructions. In brief, 10-12μm cryosections of the brain were post-fixed in 4% PFA for 10 minutes at RT. Slides were subjected to sequential dehydration in ethanol followed by Protease IV treatment for 30 minutes. Appropriate combinations of probes were prepared and added onto the slides for 2 hours at 40°C for hybridization. This was followed by four amplification steps using AMP1, AMP2, AMP3, AMP4a and counter staining for DAPI. Coverslips were mounted with Prolong Gold. Following probes were used for analysis: *Tyrobp* (ACD no. 408191-C3), *Cd9* (ACD no. 430631), *Sox10* (ACD no. 435931-C3), *Serpina3n* (ACD no. 430191). 3-4 images of the cortex were acquired for each section and TIFF images generated for separate channels were analyzed using a CellProfiler software (V2.2.0). Specifications for puncta calculation were set to a diameter of 3-12 pixels within DAPI^+^ nuclear space. For co-localization analysis, nuclei containing puncta of each probe were related using the ‘Relate objects’ function and percentage co-localization was calculated.

### Statistical analysis

Statistical analysis was performed in GraphPad Prism8 software and presented as SEM using unpaired student t-tests. Degree of significance was determined by the software and is mentioned for each data chart. For *in-vitro* experiments, technical replicates were used. For scRNAseq experiments, replicate of 3 mice per condition were used for all analysis. Immunofluorescence staining and RNAscope experiments are often performed on biological replicates and representative images are shown in the figure.

---

**Figure S1:**
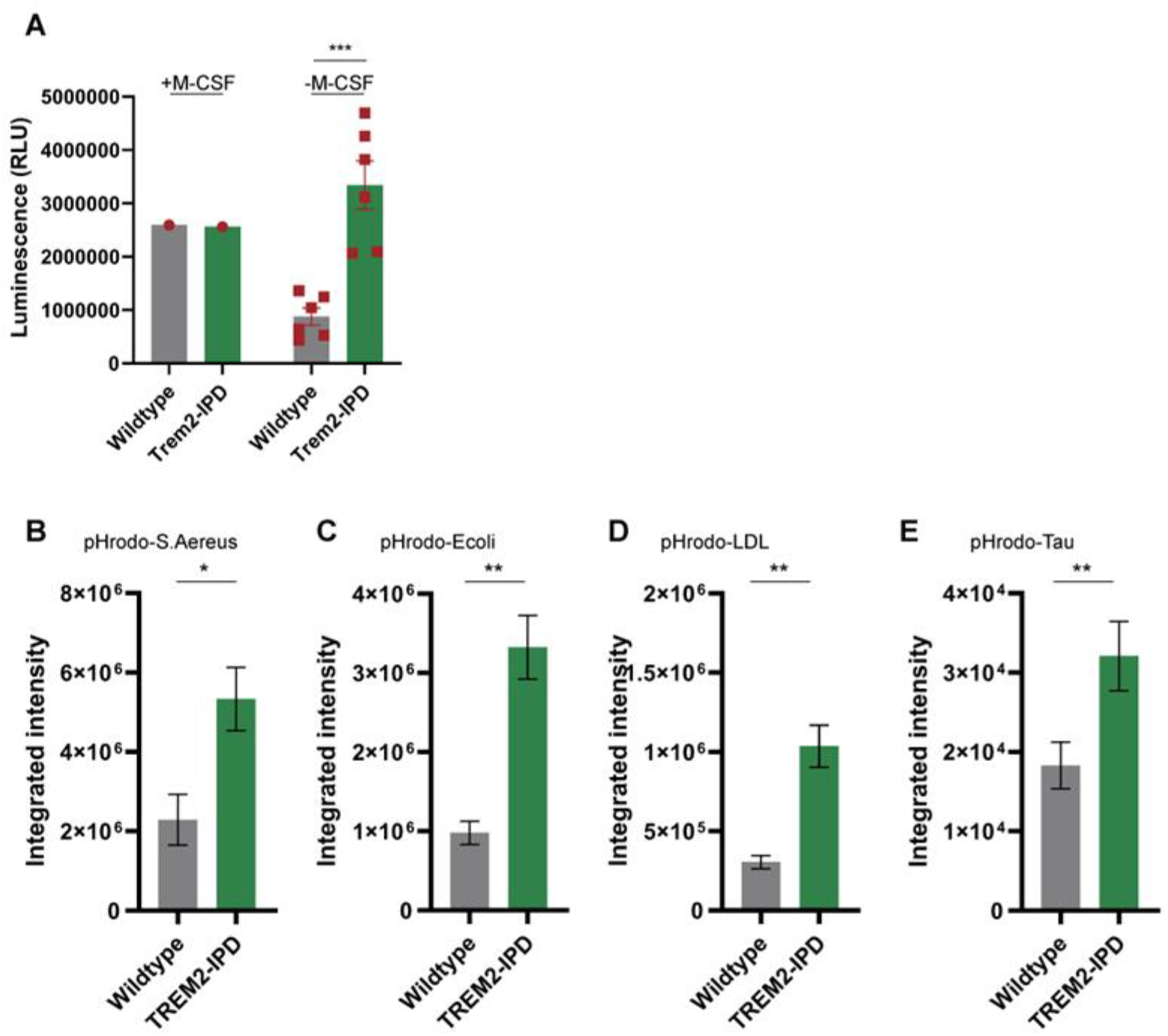
IPD substitution primes microglia activity in-vitro, related to Figure 1. (A) Cell survival assay on BMDM from WT and Trem2-IPD mice upon M-CSF withdrawal measured as levels of cellular ATP represented as luminescence (unpaired t-test; p=0.0004). *In-vitro* phagocytosis by primary microglia derived from P7 Wildtype and Trem2-IPD pups treated with (B) pHrodo-S.Aureus, (C) pHrodo-E.coli, (D) pHrodo-LDL and (E) pHrodo-Tau. *p=0.0243, **p=0.0020 Error bars represent SEM and symbols represent replicates.

**Figure S2:**
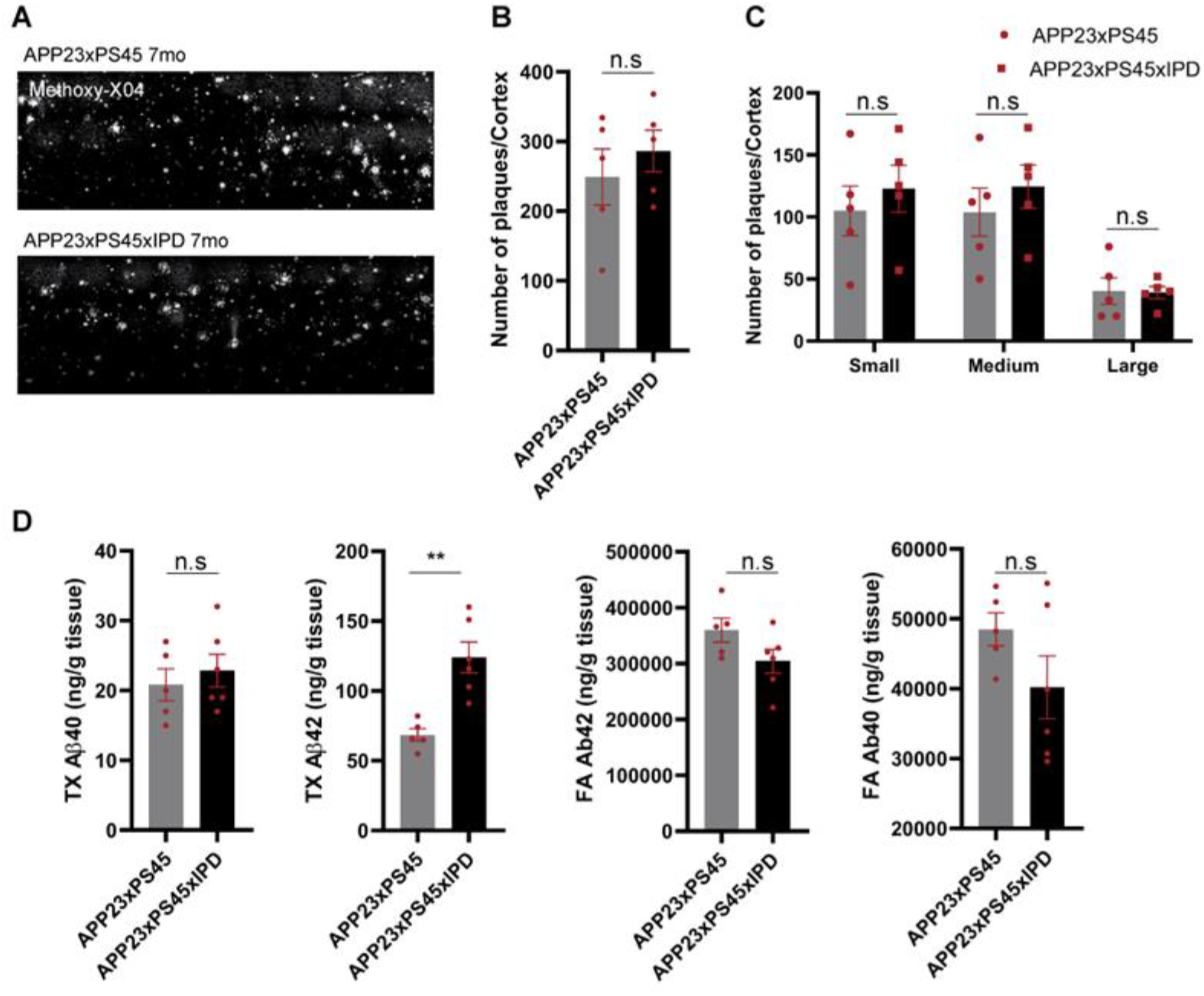
Amyloid plaque load plateaus in 7mo AD mice, related to Figure 1. (A) Representative images of Methoxy-X04 staining showing amyloid plaque load in APP23xPS45 (upper image) and APP23xPS45xIPD (lower image) mice at 7mo. (B) Quantification of number of plaques in the cortex (unpaired t-test; p=0.4809) and (C) Plaque size (unpaired t-test; small: p=0.5346, medium: p=0.4526, large: p=0.9212) in the cortex upon Methoxy-X04 staining. (D) ELISA analysis of soluble (TX fraction) Aβ40 and Aβ42 levels in forebrain lysates at 7mo (top; unpaired t-test p=0.5547 and 0.0019) and analysis of insoluble (FA fraction) Aβ40 and Aβ42 levels in forebrain lysates at 7mo (bottom; unpaired t-test; p=0.1585 and 0.1020 respectively).

**Figure S3:**
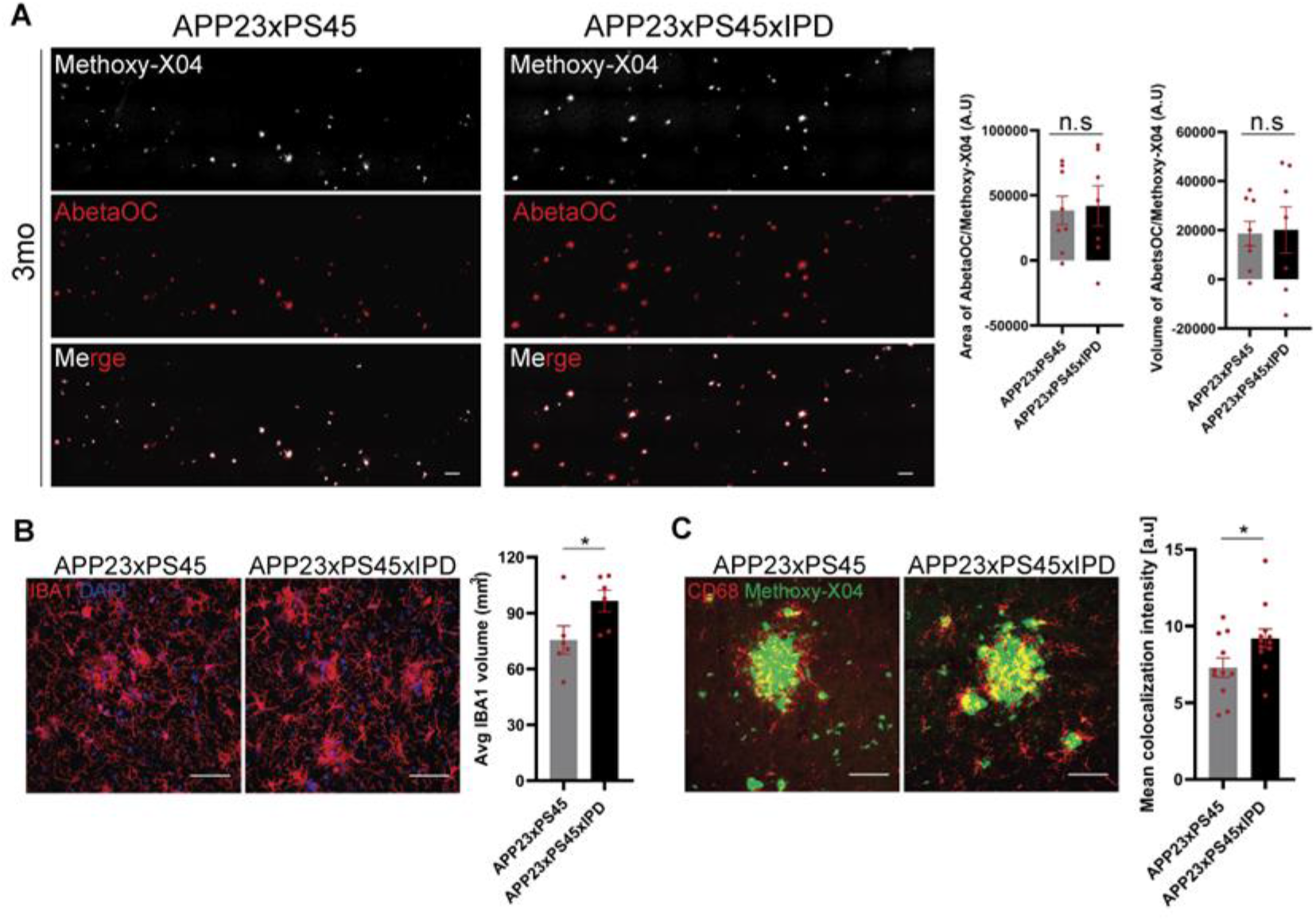
Plaque species unaffected by IPD-substitution, related to Figure 1. (A) Representative images of AbetaOC and Methoxy-X04 co-staining showing amyloid plaque load in APP23xPS45 and APP23xPS45xIPD mice at 3mo. Quantification of AbetaOC area (unpaired t-test; p=0.8543) and volume (unpaired t-test; p=0.8881) in the cortex calculated over Methoxy-X04 label. (B) Representative images of IBA1 staining and analysis of IBA1 volume per image in 3mo APP23xPS45 and APP23xPS45xIPD mice (unpaired t-test; p=0.0348). (C) Representative images of CD68/Methoxy-X04 double immunofluorescence and analysis showing increased CD68 signal intensity around Aβ plaques (unpaired t-test p=0.0319). Scale bar for A=100μm and B, C=50μm. Error bars represent SEM.

**Figure S4:**
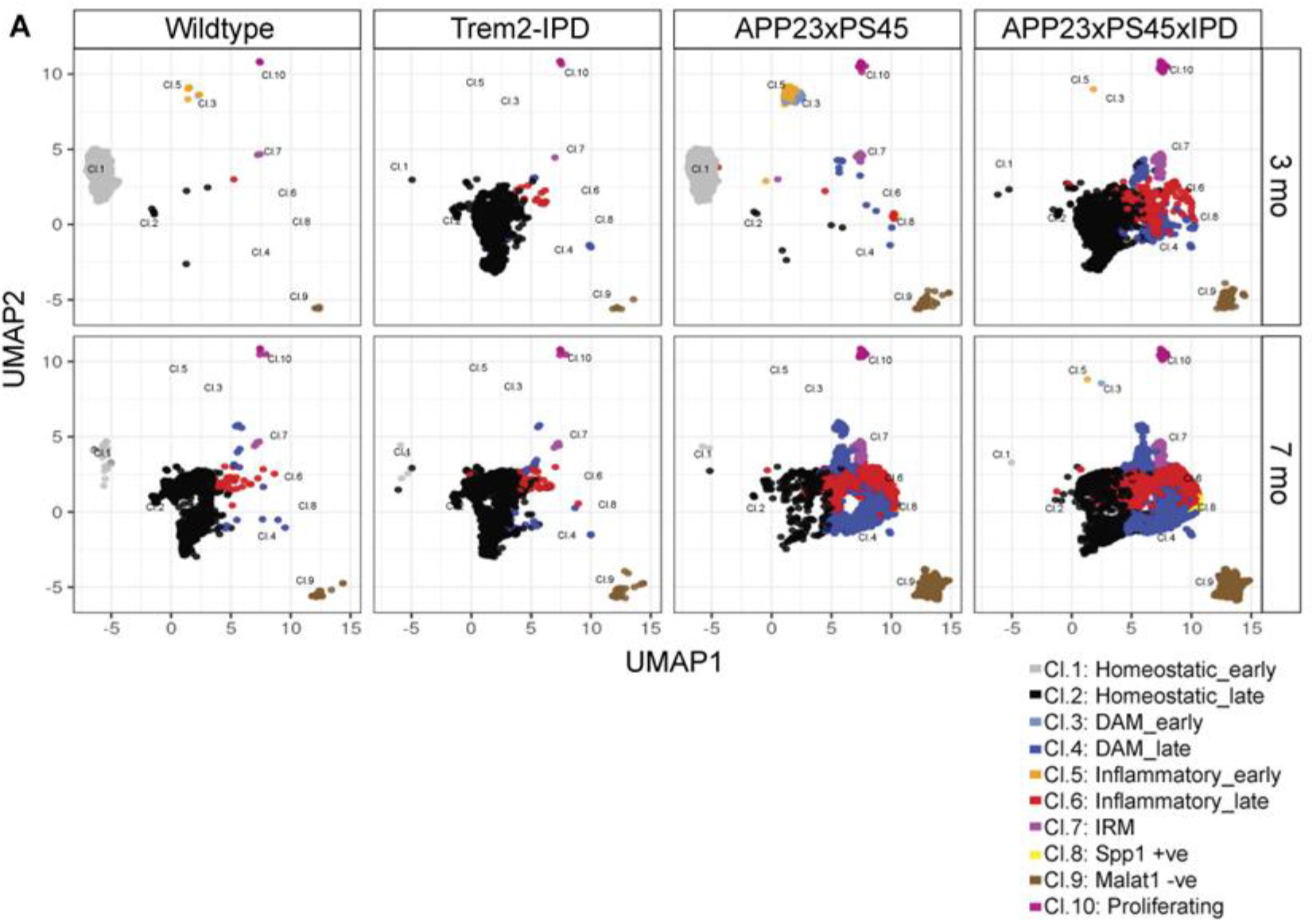
IPD-substitution shows differences in clustering between 3mo and 7mo, related to Figure3. (A) UMAP visualization of microglial clustering split by genotype and age showing an IPD-mediated shift in clusters at 3mo overlapping with 7mo clusters.

**Figure S5:**
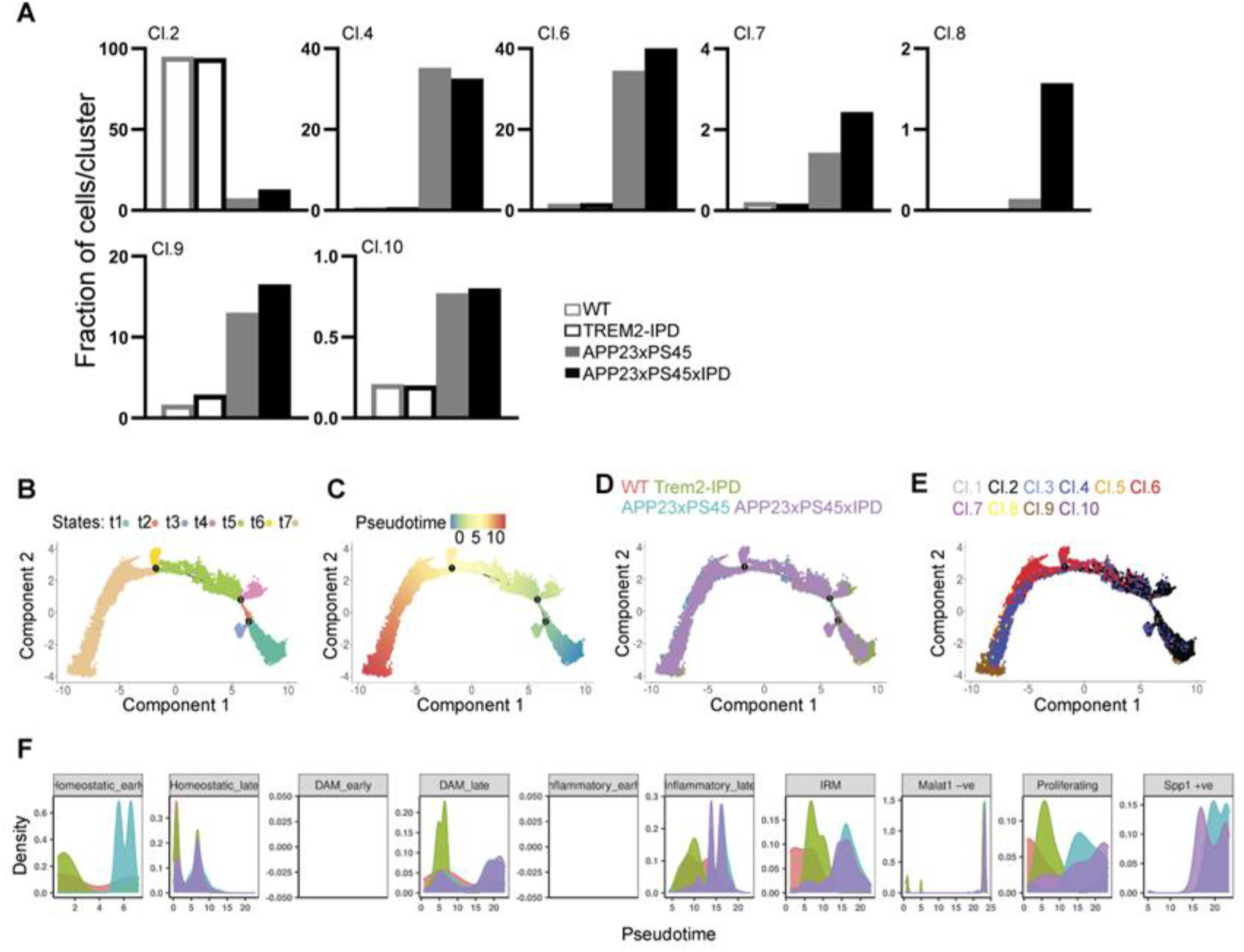
Microglial phenotype and trajectory similar in 7mo AD mice, related to Figure 4. (A) Fraction of cells belonging to each microglial cluster as labelled, displayed as bar plots from all samples at 7mo mice. Cluster 8 is composed of microglia predominantly from the IPD mice. Cell trajectory analysis of microglial sub-clusters at the 7mo time-point using Monocle based on (B) State, (C) Pseudotime (with root set to homeostatic microglia in 7mo WT mice), (D) Genotype and (E) Sub-clusters (as established in Fig.3b) showing the bifurcation of microglial progression into two separate axis driven by IPD-substitution. (F) Density distribution plot of the pseudotime showing microglia sub-clusters across all genotypes at 7mo.

**Figure S6:**
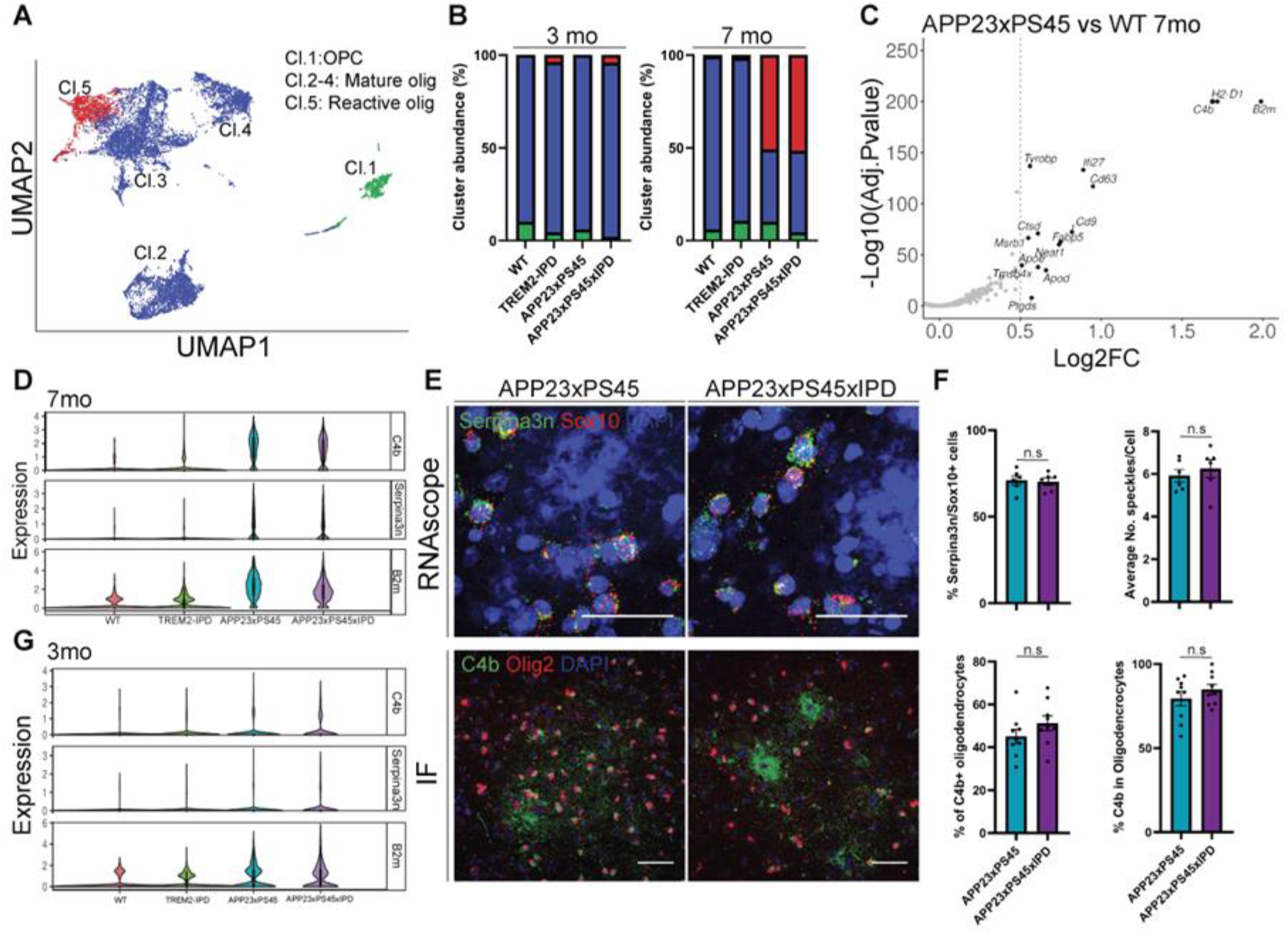
Reactive oligodendrocytes appear in AD mice and are unaffected by IPD. (A) UMAP visualization and annotation of 5 sub-clusters within the oligodendrocyte lineage across all samples; 2,300 cells at 3mo and 1,480 cells at 7mo. (B) Stacked bar graph displaying cell percentage of each cluster identifies the appearance of reactive oligodendrocytes (red bar) in 7mo APP23xPS45 and APP23xPS45xIPD mice. (C) Volcano plot showing DEGs between APP23xPS45 versus WT mice in the oligodendrocyte cluster at 7mo. (D) Violin plot showing the expression of C4b, Serpina3n and B2m within reactive oligodendrocytes (Cl.5) across all genotypes at 7mo. (E) Representative images of RNAscope-ISH for Serpina3n and Sox10 probes (upper panel) and immunofluorescence for C4b and Olig2 co-staining (lower panel) in the cortex of APP23xPS45 and APP23xPS45xIPD mice at 7mo. (F) Quantification of percentage and expression (denoted by puncta/cell) of Serpina3n and C4b in oligodendrocytes based on RNAscope-ISH and immunofluorescence images; n represents number of images from 3 independent mice. (G) Same as D, expression in 3mo mice. Error bars represent SEM. Unpaired t-test; p=0.8327, p=0.5367, p=0.3246, p=0.2215. Scale bar= 20μm upper panel, 50μm lower panel.

**Figure S7:**
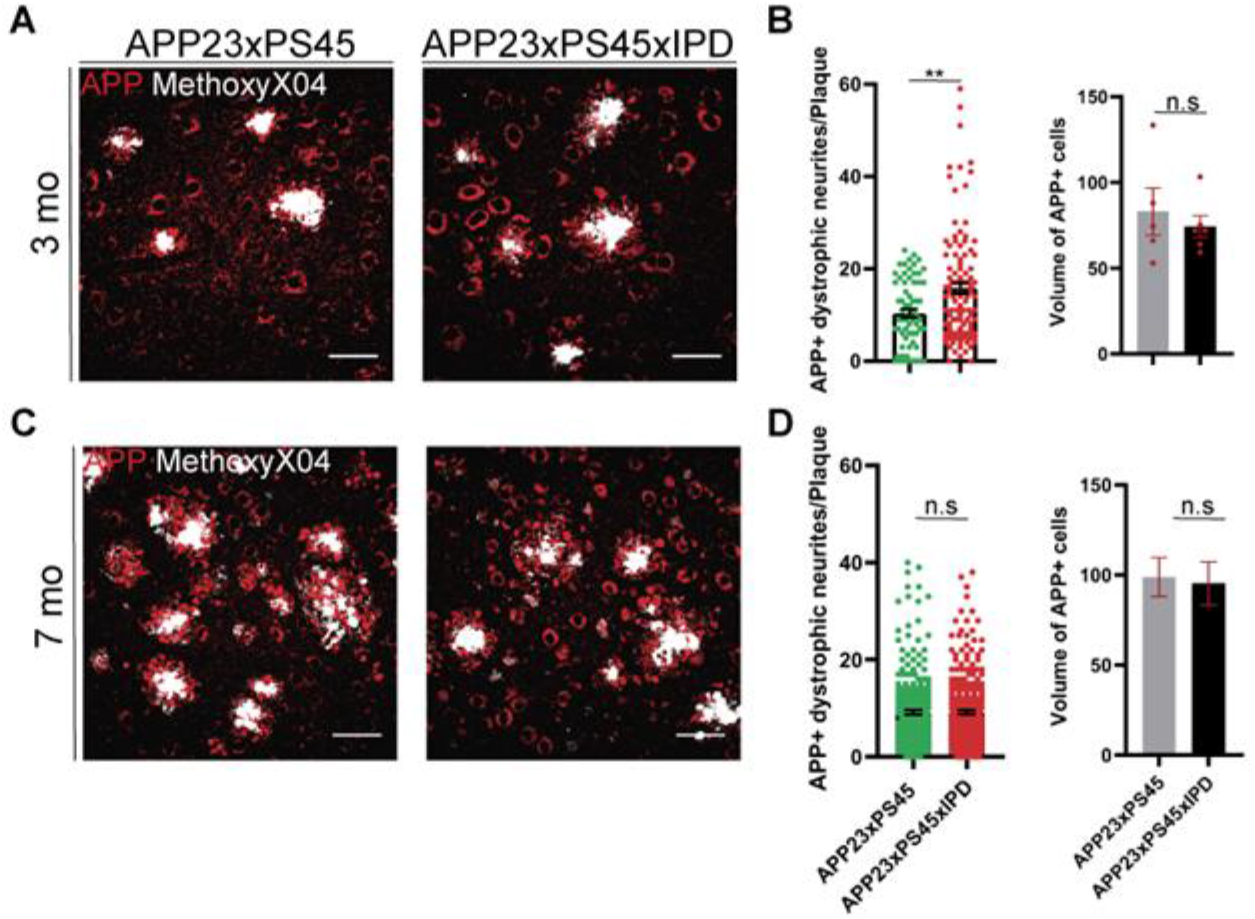
Increased neurite dystrophy upon IPD substitution at early stage pathology in AD mice. (A) Representative immunofluorescence image of APP^+^ dystrophic neurites surrounding Methoxy-X04^+^ amyloid plaques in the cortex of 3mo APP23xPS45 and APP23xPS45xIPD mice. (B) Quantification of APP^+^ neurite number (unpaired t-test; p=0.0011; symbols=number of plaques counted) and volume (unpaired t-test; p=0.55) around each plaque in 3 month old mice. (C) Same as A, but in the cortex of 7mo mice. (D) Quantification of APP^+^ neurite number (unpaired t-test; p=0.9655, symbols=number of plaques counted) and volume (unpaired t-test; p=0.8364) around each plaque in 7mo mice. Scale bar for A and C=50μm; Error bars represent SEM.

